# Distillation of MSA Embeddings to Folded Protein Structures with Graph Transformers

**DOI:** 10.1101/2021.06.02.446809

**Authors:** Allan Costa, Manvitha Ponnapati, Joseph M. Jacobson, Pranam Chatterjee

## Abstract

Determining the structure of proteins has been a long-standing goal in biology. Language models have been recently deployed to capture the evolutionary semantics of protein sequences. Enriched with multiple sequence alignments (MSA), these models can encode protein tertiary structure. In this work, we introduce an attention-based graph architecture that exploits MSA Transformer embeddings to directly produce three-dimensional folded structures from protein sequences. We envision that this pipeline will provide a basis for efficient, end-to-end protein structure prediction.

## Introduction

Elucidating protein structure is critical for understanding protein function. However, structure determination via experimental methods such as x-ray crystallography [Smyth, 2000] or cryogenic electron microscopy (cryo-EM) [Murata and Wolf, 2018], is a time-consuming, difficult, and expensive task. Classical modeling methods have attempted to solve this task *in silico*, yet are computationally prohibitive [Rohl et al., 2004, Hollingsworth and Dror, 2018, Wang et al., 2016]. Recently, machine learning approaches have been deployed to harvest available structural data and efficiently map sequence to structure [Yang et al., 2020, Senior et al., 2020].

Transformer models are sequence-to-sequence architectures that have been shown to capture the contextual semantics of words [Vaswani et al., 2017] and have been widely deployed as language models [Devlin et al., 2019, Brown et al., 2020]. The sequential structure of proteins, imposed by the central dogma, along with their hierarchical semantics, developed by Darwinian evolution, makes them a natural target for language modeling. Recently, transformers have been deployed to learn protein sequence distributions, and generate latent embeddings that grasp relevant structure [Rives et al., 2021, Elnaggar et al., 2020, Vig et al., 2020], most notably tertiary structural information [Rao et al., 2020]. Augmenting input sequences with their evolutionarily-related counterparts, in the form of a multiple sequence alignment (MSA), further strengthens the predictive power of these transformer architectures, as demonstrated by state-of-art contact prediction results [Rao et al., 2021].

In this study, we leverage MSA Transformer embeddings within a geometric deep learning architecture to directly map protein sequences to folded, three-dimensional structures. In contrast to existing architectures, we directly estimate point coordinates in a learned, canonical pose, which removes the dependency on classical methods for resolving distance maps, and enables gradient passing for downstream tasks, such as side chain prediction and protein refinement. Overall, our results provide a bridge to a complete, end-to-end folding pipeline.

## Methodology and Results

We treat the protein folding problem as a graph optimization problem. First, we harvest information-dense embeddings produced by the MSA Transformer [Rao et al., 2021], and use these embeddings to produce initial node and edge hidden representations in a complete graph. To process and structure geometric information, we employ the attention-based architecture of the Graph Transformer, as proposed by [Shi et al., 2021]. Final node representations are then projected into Cartesian coordinates through a learnable transformation, and the resulting induced distance maps are compared to ground truth to define the loss for training.

### MSA Transformer Data Augmentation

The MSA Transformer is an unsupervised protein language model that produces information-rich residue embeddings [Rao et al., 2021]. In contrast to other protein language models, it operates on two dimensional inputs consisting of a length-*N* query sequence along with its MSA sequences. It utilizes an Axial Transformer [Ho et al., 2019] as an efficient attention-based architecture for performing computation on its layers’ *O*(*N · S*) representations, where *S* is the total number of input MSA sequences.

Our model seeks to operate on graph features distilled from MSA Transformer encodings. Last-layer residue embeddings capture individual and contextual residue properties. Similarly, the vector formed by pairwise attention scores at each layer and head captures attentive interactions between residue pairs. The richness of information present at these vectors was demonstrated in state-of-the-art contact prediction [Rao et al., 2021]. We perform the natural step of extending those individual and pairwise embeddings to node and edge representations, and show how learning over the resulting graph can resolve a protein’s three-dimensional structure.

For this study, we employed the 100 million parameter-sized ESM-MSA-1 model [Rao et al., 2021], which was trained on 26 million MSAs queried from UniRef50 and sourced from UniClust30. ESM-MSA-1 produces *N* residue embeddings 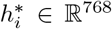 and *N × N* attention score traces 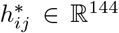 for each input sequence. Since the MSA Transformer is computationally expensive to evaluate for large *S* even in the context of inference, we precomputed the encodings and made them readily available for training. We used *S* = 64 and stored residue embeddings 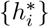 and attention score traces, 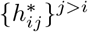 for each query sequence.

For training and validation, we used the ESM Structural Split [Rives et al., 2021], which builds upon trRosetta’s training dataset [Yang et al., 2020]. To overcome the bottleneck associated with reading large encodings directly from the file system, we fixed our splits to the first superfamily split as specified in Rives, et al., and serialized its MSA Transformer encodings into tar shards. A virtual layer of data shuffling was added through the WebDataset framework [Aizman et al., 2020]. The resulting dataset of graph features has 0.25 TB.

## Graph Building

We treat a protein as an attributed complete graph. Let *H*_*V*_ and *H*_*E*_ be the dimensionalities of node and edge representations, respectively. These attributes are extracted from MSA-Transformer embeddings through standard deep neural networks:

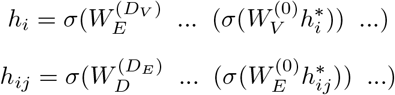

Where 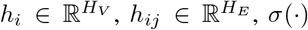 is a ReLU nonlinearity, and *D*_*V*_ and *D*_*E*_ are the depths of node and edge information extractors, respectively. *W* denotes dense learnable parameters, and here and in the following equations we omit bias terms.

## Graph Transformer

The Graph Transformer was introduced in [Shi et al., 2021] to incorporate edge features directly into graph attention. This is possible by directly summing transformations of edge attributes to the original keys and values of the attention mechanism. We approach protein folding with a variation of this architecture. Consider layer *l* node hidden states, 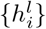, and similarly learned edge latent states {*e*_*ij*_}. If we employ *C* attention heads, a layer update can be written as

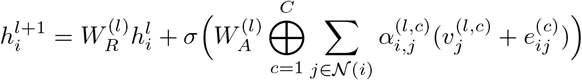

Where ⊕ denotes concatenation, 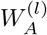 and 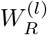 are learnable projections. As in the original architecture, we apply batch normalization to each layer. The attention scores 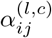, node values 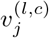 and edge values 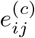 are obtained from learnable transformations of the original node hidden states and edge attributes:

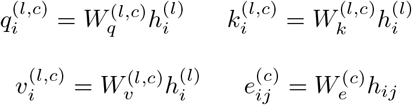

The attention scores are normalized according to graph attention:

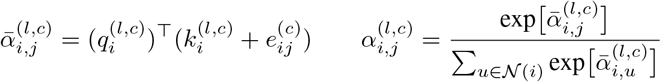

To hold computational costs roughly constant, we let 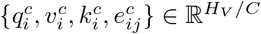 as in standard Transformer architectures.

### Cartesian Projection and Loss

We train a predictor to recover coordinates of each residue in a learned canonical pose:

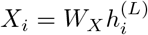

Where *X*_*i*_ ∈ ℝ^3^. To train our network, we use a distogram-based loss function on the resulting distance map. Let 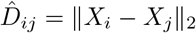 be the induced Euclidean distance between the Cartesian projections of nodes *i* and *j*, and *D*_*ij*_ be the ground truth distance. Our loss is based on the *L*_1_-norm of the difference between those values:

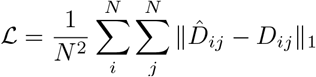

### Model Training

To optimize our trained model, we performed a shallow random hyperparameter search for *H*_*V*_ ∈ {32, 64, 128, 256}, *H*_*E*_ ∈ {32, 64, 128}, *L* ∈ {3, 6, 10, 15}, *C* ∈ {1, 2, 4}. We utilized the Adam Optimizer with lr ∈ {1 × 10^−3^, 3 × 10^−4^, 1 × 10^−4^, 3 × 10^−5^, 1 × 10^−5^}. We further tested variations of the loss function, testing the MSE loss and weighted versions of *L*_1_ and MSE for batch sizes *B* ∈ {10, 15, 30}.

To handle GPU memory constraints, we employed gradient checkpointing at each Graph Transformer layer. Models were trained in parallel on NVIDIA V100s provided by the MIT SuperCloud HPC [Reuther et al., 2018].

In total, we performed 40 search training runs, with a maximum of 70 epochs and an early stop with a patience of 3 for validation loss. Our best model trained for 17 hours without registering early stopping. With *H*_*V*_ = *H*_*E*_ = 64, *L* = 10 and *C* = 1, this model only possesses a total of 382K parameters. We used lr= 3 × 10^4^ and *B* = 30, as well as an *L*_1_ loss, and achieved ℒ_val_ = 2.25 and GDT_TS_val_ = 40.58.

### CASP13 Evaluation

To investigate the generalization of our model, we evaluated it on the free modeling targets from the 13th edition of the Critical Assessment of Protein Structure Prediction (CASP13). We chose to benchmark our model against the performance of the current state-of-the-art public architecture: trRosetta [Yang et al., 2020]. trRosetta considers a sequence’s MSA to predict distance probability volumes as well as relevant interresidue orientations. In contrast to our model, trRosetta relies on restraints derived from the predicted distance and orientations for downstream Rosetta minimization protocols [Rohl et al., 2004]. For each distance, we consider trRosetta’s best prediction as its expected value or its maximum likelihood estimate. We utilized dRMSD (distogram RMSD) between predicted distances and ground truth as our evaluation metric. To make a direct comparison, we only consider distances that lie within trRosetta’s binning range (2-20 Å).

Our results demonstrate that the Graph Transformer model, despite its size, is competitive to trRosetta’s estimates (Figure 3, Table 1). It is worth noting that our architecture resolves backbone structure as its main output and uniquely and deterministically produces distances, whereas trRosetta operates within a probabilistic domain that does not need three-dimensional resolution. These early results thus suggest potential for improved predictive capability with larger model capacity and downstream protein refinement.

**Table 1:**
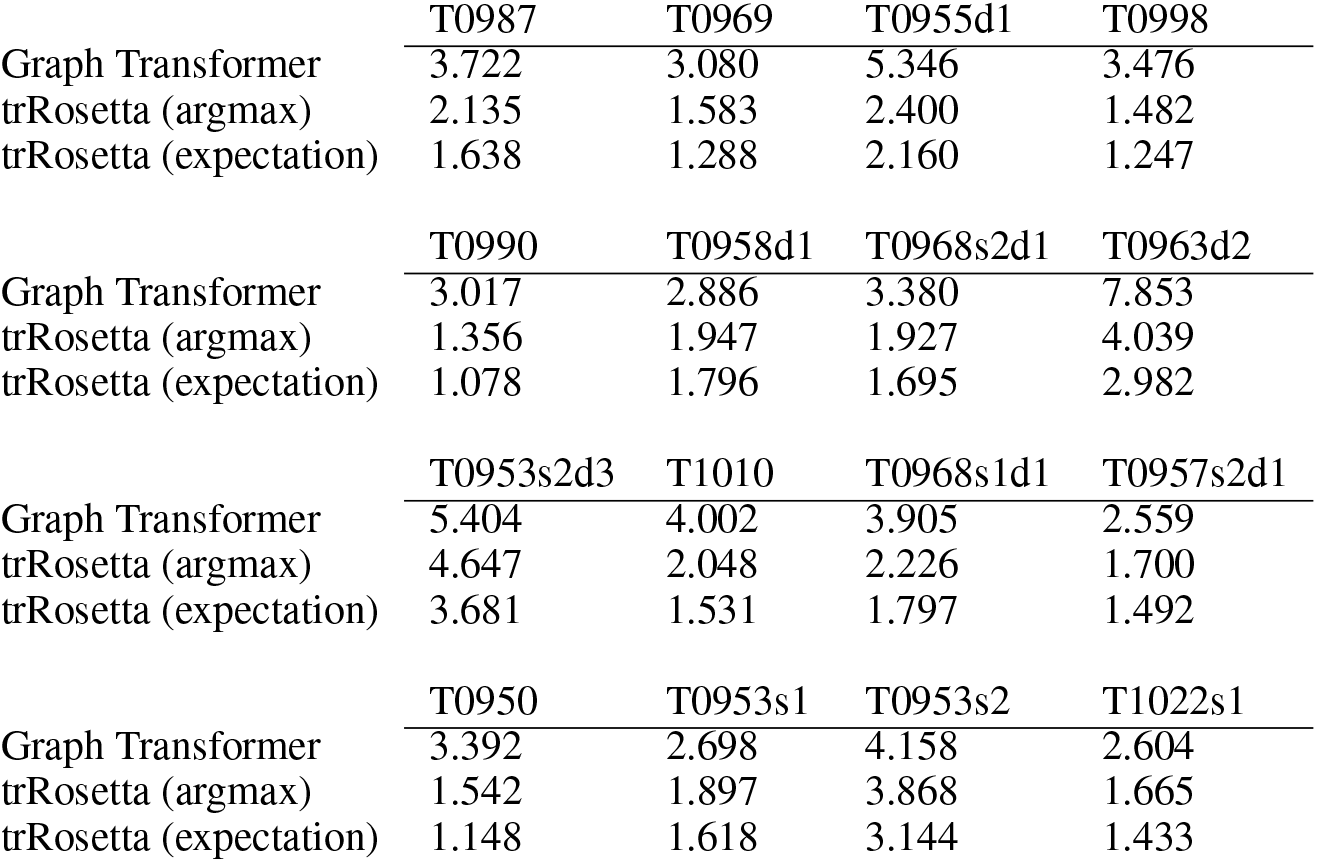
CASP13 Free Modeling benchmarks of dRMSD for our architecture’s induced distances and trRosetta’s expectation and argmax distances, against ground truth. We only consider distances that lie within trRosetta’s binning range.

**Figure 1:**
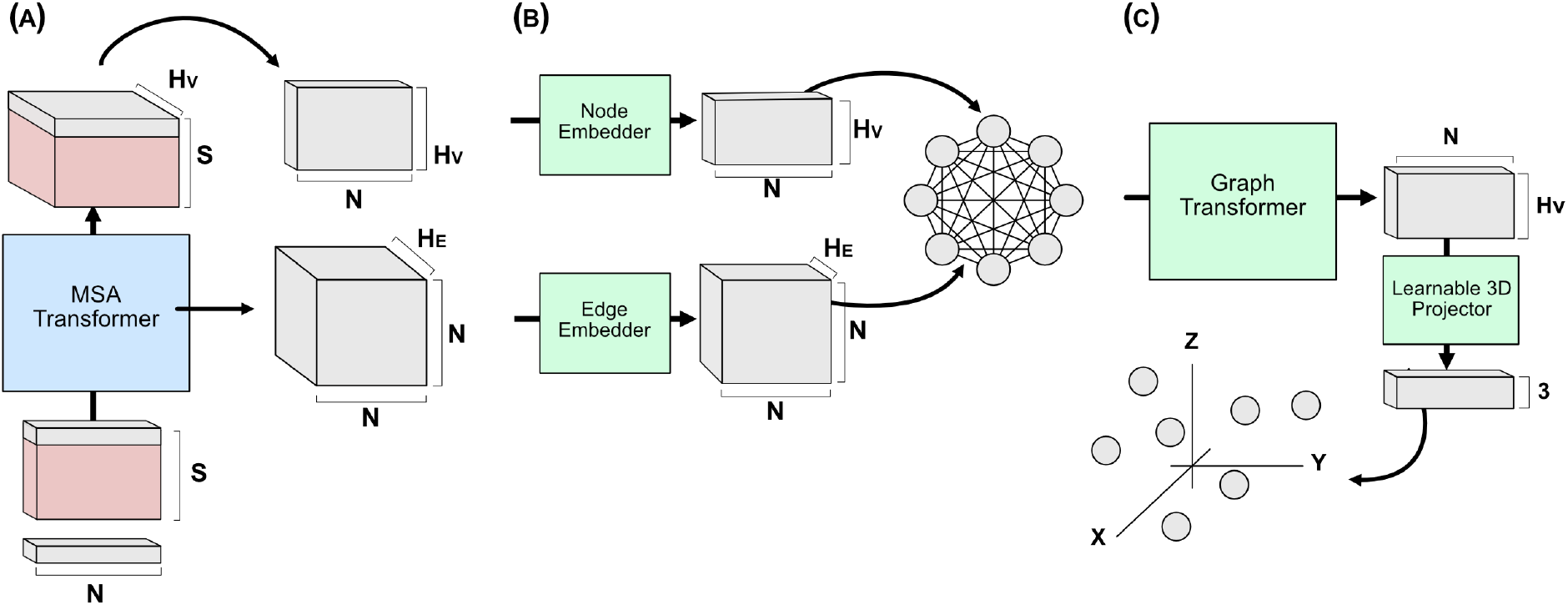
An overview of a sequence-to-structure pipeline utilizing the MSA-Transformer and a Graph Transformer. **(a)** We first augment a length-*N* protein sequences to *S* of its MSA. The MSA-Transformer operates over this token matrix to produce enriched individual and pairwise embeddings. We store those embeddings that are from the original query sequence. **(b)** Deep neural networks extract relevant features and structure latent states for a downstream graph transformer. Individual and pairwise embeddings are assigned to nodes and edges, respectively. **(c)** A graph transformer operates on node representations through an attention-based mechanism that considers pairwise edge attributes. The final node encodings are projected directly to ℝ^3^, and the induced distogram is computed for the loss.

**Figure 2:**
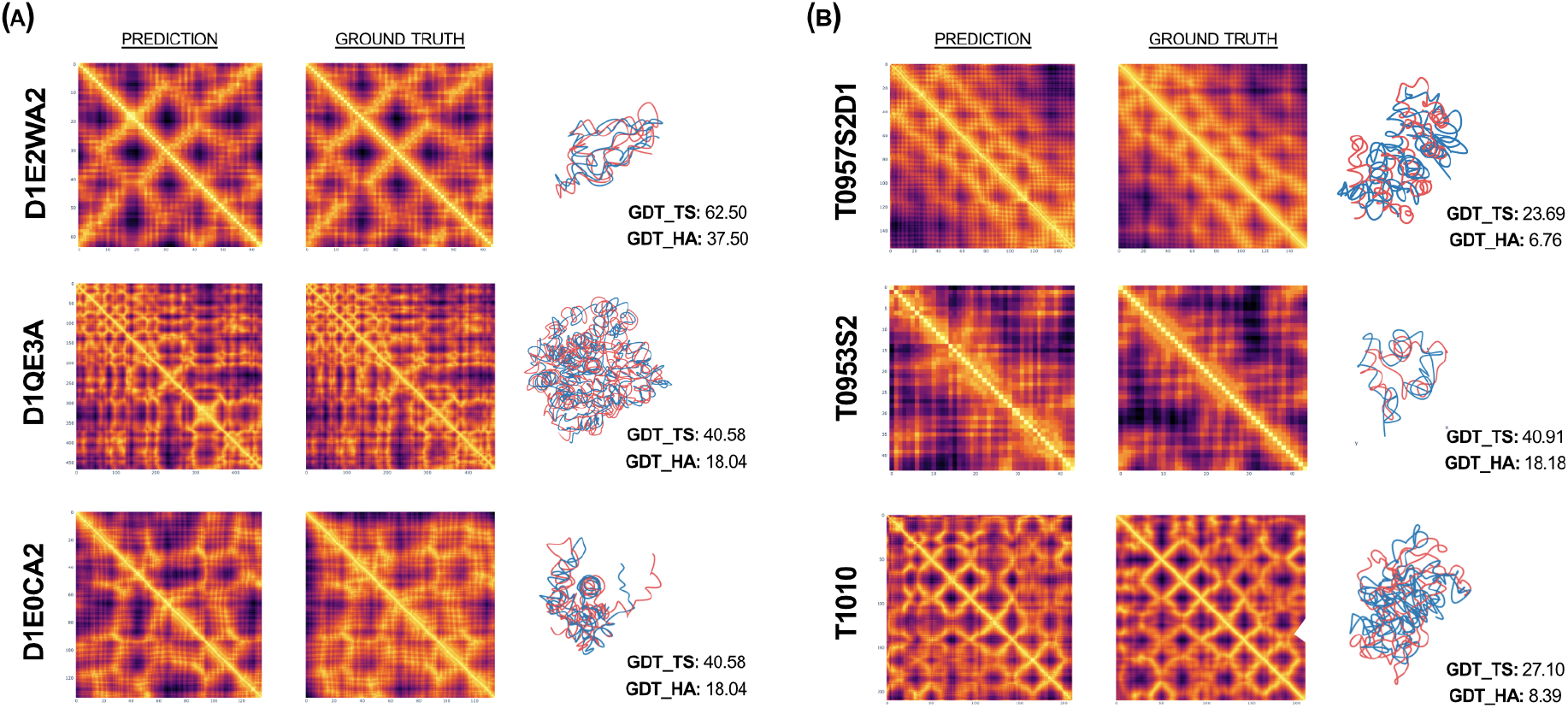
Comparison of predicted distograms and three-dimensional arrangement with their ground truth counterparts for samples from (A) ESM Structural Split dataset, and (B) CASP13 free modeling targets. For each prediction-ground truth pair, a PDB name is indicated on the left. Aligned *Cα* traces and metrics are indicated on the right. Blue traces denote model predictions, red traces denote ground truth. Traces were produced by fitting splines to the sequence of predicted *Cα* coordinates.

**Figure 3:**
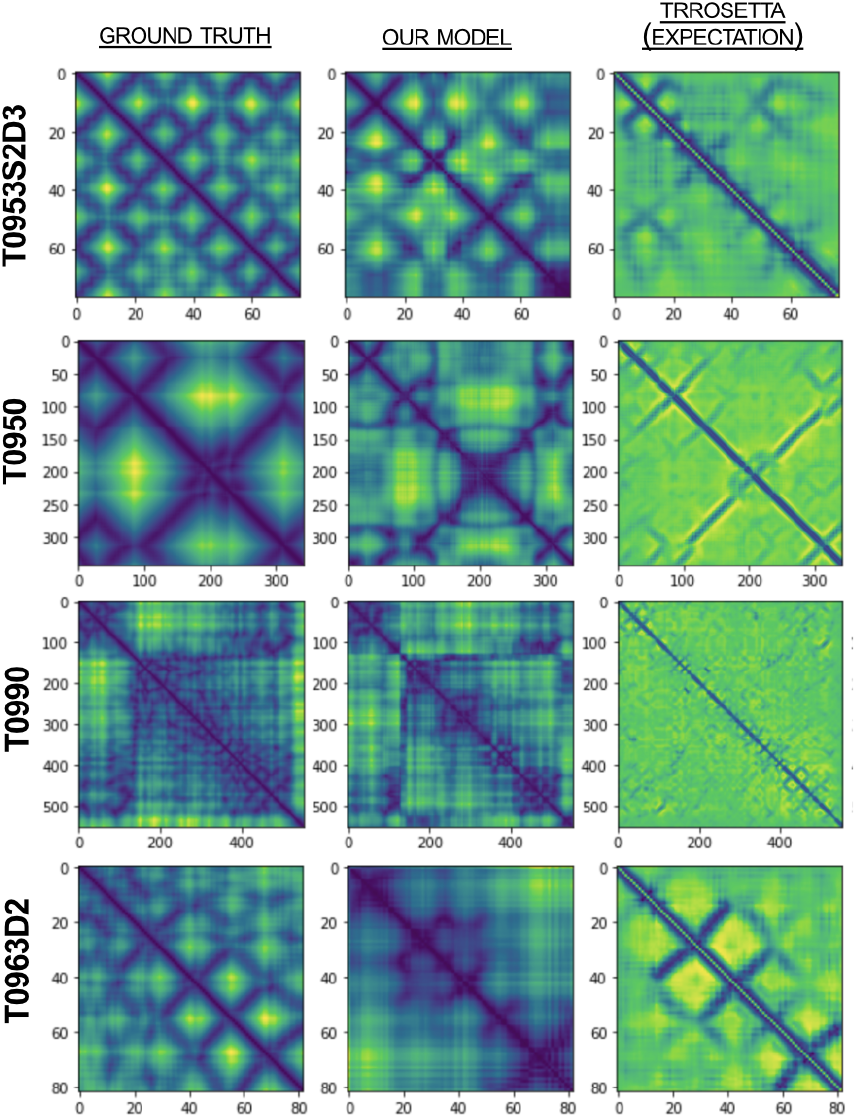
Qualitative assessment of model predictions for CASP13 free modeling targets. Note that our model is able to capture long range interactions, whereas trRosetta by construction is limited to short range dependencies. We highlight T0950 an T0963D2 as examples of challenging reconstructions for our network.

## Discussion

In this work, we revisit the protein folding problem and highlight the role of unsupervised language models in providing a meaningful basis for the sequence-to-structure prediction task. We present a strategy to encapsulate MSA Transformer embeddings and attention traces in a geometric framework, and formalize a graph learning pipeline to reason positional information.

Overall, our results demonstrate the remarkably expressive power of language models and, in particular, of MSA-augmented architectures. To demonstrate a versatile bridge between sequence and three-dimensional structure, we trained a downstream model to produce *Cα*-traces which, before any refinement is performed, induce distograms with high similarity to ground truth.

We emphasize that our model, in its current state, tackles only a step of the protein structure prediction problem. With only 382K parameters, it serves as a fast and scalable solution to resolving the position of protein backbones. Furthermore, it extends learning beyond distogram prediction and provides a natural foundation for downstream tasks, such as side chain prediction and protein refinement. Most importantly, we hypothesize that by increasing model capacity, dataset size, and training time, our model’s predictive capability can improve significantly.

In total, this study builds upon the recent groundbreaking work in protein representation learning and protein language modeling by the community. The integration of diverse network architectures and pretrained models, as demonstrated here, will enable the eventual efficient solution of the protein structure prediction problem.

## Acknowledgements

We thank the MIT SuperCloud and Lincoln Laboratory Supercomputing Center for providing the HPC and database resources that have contributed to the research reported within this manuscript. We thank Thrasyvoulos Karydis for the fruitful discussions and insights for this project.

